# Theta and alpha oscillations are traveling waves in the human neocortex

**DOI:** 10.1101/218198

**Authors:** Honghui Zhang, Andrew J. Watrous, Ansh Patel, Joshua Jacobs

**Affiliations:** School of Biomedical Engineering, Columbia University, New York, NY 10027

## Abstract

Human cognition requires the coordination of neural activity across widespread brain networks. Here we describe a new mechanism for large-scale coordination in the human brain: traveling waves of theta and alpha oscillations. Examining direct brain recordings from neurosurgical patients performing a memory task, we found contiguous clusters of cortex in individual patients with oscillations at specific frequencies between 2 to 15 Hz. These clusters displayed spatial phase gradients, indicating that the oscillations were traveling waves that propagated across the cortex at ∼0.25-0.75 m/s. Traveling waves were relevant behaviorally because their propagation correlated with task events and was more consistent when subjects performed the task well. Our findings suggest that traveling waves can be modeled by a network of coupled oscillators because the direction of wave propagation correlated with the spatial orientation of local frequency gradients. These findings suggest a role for traveling waves in supporting brain connectivity by organizing neural processes across space and time.

## Introduction

Oscillations have a distinctive role in brain function because they coordinate neuronal activity on multiple scales. Brain oscillations are important at the microscale, because they modulate the timing of neuronal spiking (Bragin et al., 1995; Jacobs et al., 2007), and at the macroscale, where they synchronize distributed cortical networks that are communicating (Fries, 2005). Owing to oscillations’ ability to coordinate neural processes across multiple scales, characterizing their spatiotemporal properties may reveal how neurons across multiple regions are dynamically coordinated to support behavior (Kopell et al., 2014).

The human cortex displays oscillations at various frequencies during cognition (Buzsaki and Draguhn, 2004). To understand how these patterns relate to behavior, researchers have generally examined the properties of oscillations at individual frequencies in local networks (Raghavachari et al., 2006; Jacobs et al., 2007) or in point-to-point links between distinct regions (Watrous et al., 2013). These approaches ignore a key feature of cortical oscillations that emerged from animal studies—that oscillations at multiple frequencies form spatially continuous neural patterns (Freeman and Schneider, 1982; Freeman et al., 2000; Agarwal et al., 2014).

One such pattern is a traveling wave, which consists of a spatially coherent oscillation that moves progressively across the cortex, appearing like a wave moving across water. Traveling waves have been studied most extensively in animal models, where they are observed most often in fine-scale recordings and were shown to be functionally important to various behaviors, including visual perception (Zanos et al., 2015), spatial navigation (Lubenov and Siapas, 2009; Agarwal et al., 2014), and movement (Rubino et al., 2006). In conjunction with predictions of computational models, these studies suggest that traveling waves are a key mechanism for guiding the spatial propagation of neural activity and computational processes across the brain (Ermentrout and Kleinfeld, 2001).

Given the potential importance of spatially coordinated brain oscillations for large-scale cortical processes, several studies tested for large-scale synchronized oscillations—a prerequisite for traveling waves—in the human cortex. However, in humans such synchrony was rare or present only on a small scale (Bullock et al., 1995; Menon et al., 1996; Raghavachari et al., 2006). These results shed doubt on the possibility that spatially coordinated oscillations such as traveling waves figured prominently in large-scale cortical processing in humans.

We re-examined the potential role of cortical traveling waves in human cognition by analyzing electrocorticographic brain (ECoG) recordings from seventy-seven neurosurgical patients (Table S1). We analyzed the data with a new analysis technique that identifies traveling waves at the single-trial level across various frequencies and electrode configurations. As we describe below, we found alpha and theta traveling waves in 96% of subjects. Traveling waves were present across a wide frequency range (2 to 15 Hz), and were relevant behaviorally, as their propagation correlated with events and subject performance in a memory task. Our results suggest that human behavior is supported by spatial patterns of theta- and alpha-band oscillations that propagate across the cortex.

## Results

To identify traveling waves in the human cortex, we examined direct electrocorticographic (ECoG) brain recordings from neurosurgical patients performing a working-memory task (Sternberg, 1966). This task was shown previously to elicit large-amplitude oscillations related to memory at various frequencies (Raghavachari et al., 2001; Jacobs and Kahana, 2009). Here, we analyzed these data using a new analytical framework that can identify synchrony and traveling waves by characterizing the spatiotemporal structure of the oscillations in each patient individually.

### Spatial frequency clustering of oscillations

A traveling wave in the cortex of one patient would appear in ECoG signals as a neuronal oscillation that is visible simultaneously on multiple electrodes at the same frequency with a systematic timing (or phase) gradient across space, such that the oscillation appears to be propagating across space (Ermentrout and Kleinfeld, 2001). Thus, a first requirement for neural oscillations across multiple electrodes to display a traveling wave is that the signals must exhibit oscillations at the same frequency. Our first step in identifying human cortical traveling waves was to find any clusters of cortex where multiple neighboring electrodes recorded oscillations at the same frequency. To identify these patterns we examined the recording from each electrode individually and measured the frequencies of any narrowband oscillations that were present in the power spectrum. We distinguished these narrowband oscillations by using a peak-picking algorithm, which found the narrowband oscillatory peaks that were elevated over the background 1/*f* ECoG power spectrum (Manning et al., 2009).

Using this technique, we identified electrodes with narrowband oscillations at various frequencies (Fig. S1A-D). In most patients there were spatially contiguous clusters of electrodes with narrowband oscillations at the same frequency (Fig. S1E-G). We refer to a contiguous group of four or more electrodes with oscillations at the same frequency as an *oscillation cluster* (see *Methods*). Across 77 patients, we found a total of 208 oscillation clusters. Oscillation clusters were present at frequencies from 2 to 15 Hz, involved 58% of all electrodes, and were present in 96% of patients.

The frequencies of oscillation clusters often differed across individuals even for electrodes in the same anatomical region (Fig. S2). This suggested to us that oscillation clusters reflected distinctive cortical networks that were individualized for a given patient. To assess whether there were true intersubject differences in the frequencies of oscillation clusters, we tested for a spatial autocorrelation in the frequencies of narrowband oscillatory peaks across the electrodes in each subject using Moran’s *I* statistic (Moran, 1950). This analysis tested the hypothesis that the frequencies of oscillations were more correlated between neighboring electrodes in a patient compared to electrodes at similar anatomical locations in other patients. We thus computed *I* for each subject and compared the results with those from a shuffling procedure that randomly interchanged electrodes between subjects. The mean within-subject frequency correlations (*I* = 0.03) were higher in the real data than in the shuffled data (*p* < 10^−3^; Fig. S1G). This result indicated that clusters of electrodes with narrowband oscillations at the same frequency reflected robust within-subject spatial frequency clustering.

The widespread presence of oscillation clusters indicates that neuronal oscillations at a single frequency were present across large regions of the human cortex. If the timing of these oscillations were synchronized, it could provide evidence for large-scale oscillatory networks (Kopell et al., 2014). Thus, we next characterized the timing of activity across each oscillation clusters to identify patterns of phase synchrony such as traveling waves (Prechtl et al., 1997; Rubino et al., 2006; Patel et al., 2012; Patten et al., 2012; Bahramisharif et al., 2013; Zhang and Jacobs, 2015).

### Identifying human cortical traveling waves

Visual inspection of the signals from many oscillation clusters indicated that their phases were synchronized and that they indeed contained traveling waves. As one example, Figure 1A–C shows the activity across an 8-Hz oscillation cluster in an electrode grid from Patient 1. While 8-Hz oscillations were visible on all channels in this grid, the relative timing of this signal varied systematically with electrode location, such that the onset time of each oscillation correlated with the electrode’s anterior–posterior position.

**Figure 1:**
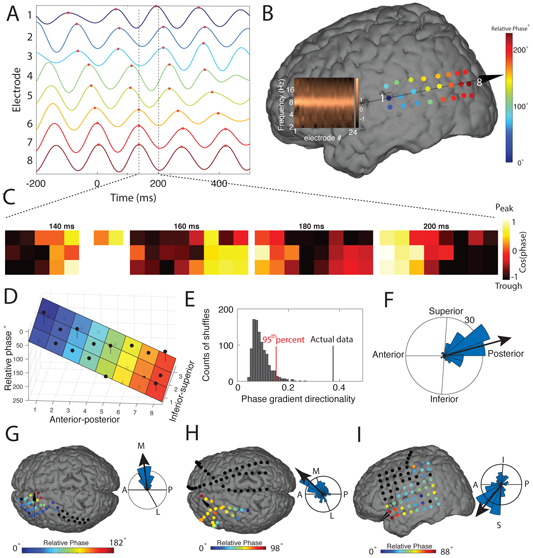
Evidence for traveling waves in the human neocortex. (A) An 8.3-Hz traveling wave in Patient 1. The plot shows this wave in one trial on electrodes ordered from anterior (top) to posterior (bottom). Peaks of individual oscillations are labeled with red asterisks, illustrating an increasing phase shift from anterior to posterior. (B) Relative phase of this traveling wave across the 3x8 electrode grid. Each electrode's color indicates the relative phase. Inset shows the normalized power spectrum for each electrode, demonstrating that all electrodes in the grid exhibit narrowband 8.3-Hz oscillations. Arrow indicates direction of wave propagation. (C) The temporal evolution of the phase pattern for the trial from Panels A & B. (D) Illustration of the fitted circular-linear model for quantifying spatial phase gradients and identifying traveling waves, applied to data from Panels A–C. Black dots indicate the average relative phase for each electrode; colored surface indicates the fitted phase plane; black lines indicate residuals. (E) Median phase gradient directionality (PGD) for this wave (black line). Gray bars indicate the distribution expected by chance. (F) Histogram indicating the directions of traveling-wave propagation observed on these electrodes across trials. (G-I) Phase distribution and direction for example traveling waves from Patients 3 (G), 63 (H), & 77 (i).

We quantified this phenomenon by calculating the relative phase of the oscillation on each electrode and trial (Fig. 1B). On this trial, oscillations showed a progressive spatial phase shift of ∼240°. In this scheme, positive phase shifts correspond to oscillators that are further advanced. Thus, because the phase was largest at posterior electrodes, it indicates that on this trial there was an anterior-to-posterior traveling wave (see *Supplemental Movie* S1).

We used circular statistics (Fisher, 1993) to measure the presence and properties of human cortical traveling waves by identifying patterns of oscillatory phase that shift systematically across the cortex. For each oscillation cluster, at each timepoint we used a circular–linear model to characterize the relation between electrode position and oscillation phase (Fig. 1D). This procedure models each cluster’s instantaneous phase distribution as a planar wave, finding the best fitting spatial phase gradient. The fitted phase gradient provides a quantitative estimate of the speed and direction of traveling wave propagation (Fig. S3). We compute the quality of fit for each traveling wave on an electrode cluster by computing the proportion of phase variation that is explained by the planar wave model, which we call the phase gradient directionality (PGD).

We assessed whether each oscillation cluster exhibited a reliable traveling wave by testing its PGD value with a permutation procedure (Fig. 1E). This analysis demonstrated that traveling waves on the cluster in Figure 1A–C were statistically reliable (mean *PGD* = 0.35, *p* < 0.001) and consistently moving in an anterior-to-posterior direction (*r*̅ = 0.94, Rayleigh *p* < 0.001; Fig. 1F). Figure 1G–I shows example traveling waves in other patients that were also robust statistically.

We applied this methodology across our dataset and found that 140 (67%) of all oscillation clusters had consistent traveling waves, thus showing both reliable planar waves at the single trial level and a reliable direction of propagation. Traveling waves involved 47% of all electrodes and were present in all lobes of the neocortex across both left and right hemispheres (Table S2). Thus, traveling waves are a broad phenomenon across the human brain.

To examine regional differences in traveling waves, we compared the properties of oscillation clusters identified from different brain areas (Fig. 2A; *Supplemental Movie* S2). Traveling waves from electrodes in the frontal and temporal lobes generally propagated in a posterior-to-anterior direction (*p*’s < 0.01, Rayleigh tests; Fig. 2B). In the occipital and parietal lobes, traveling waves had a range of propagation directions that were not reliably clustered (*p* > 0.05). In the frontal lobe, traveling waves often propagated towards the midline.

**Figure 2:**
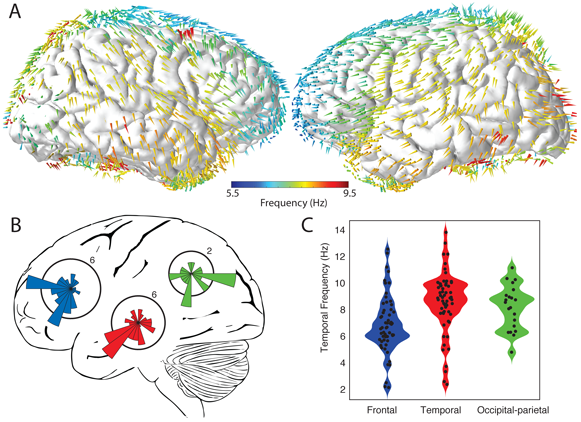
Population analysis of traveling wave direction and frequency. (A) Spatial topography of mean traveling-wave direction and frequency. Colored arrows indicate the mean direction and frequency of traveling waves observed at an electrode within 1.5 cm. (B) Distribution of the mean direction of traveling waves from each lobe. The orientations of the polar histograms are projected to match the lateral brain view. (C) Distributions of temporal frequencies for traveling waves from different regions. Black dots indicate the mean frequency from individual electrode clusters.

We also compared the temporal frequencies of the oscillations that were involved in traveling waves (Fig. 2C). Individual traveling waves were present at frequencies ranging from 2 to 15 Hz. Traveling waves in the frontal lobe had slower mean temporal frequencies in the “theta” range (mean 6 Hz). In contrast, traveling waves in occipital and temporal regions had faster “alpha”-like frequencies (mean 9 Hz; ANOVA, *F*_(2,137)_ = 9.7, *p* < 0.01). It is notable that the frequencies of the traveling waves we found in frontal and occipital regions were similar to the theta and alpha oscillations that are commonly measured in these areas (Klimesch, 1999; Voytek et al., 2010; Groppe et al., 2013) because it suggests that these other oscillations could be traveling waves.

### Behavioral relevance of cortical traveling waves

We hypothesized that the spatial propagation of traveling waves reflected the movement of neural activity across the cortex in a manner that was important for behavior. Although some previous studies had measured human cortical traveling waves during tasks, they did not show clear correlations to features of behavior (Massimini et al., 2004; Takahashi et al., 2011; Bahramisharif et al., 2013). We tested for a potential functional role for traveling waves by comparing their properties through the course of memory processing. In each trial of this memory task (Sternberg, 1966), patients learned a list of stimuli and then viewed a retrieval cue. By comparing traveling-wave properties during this period, we hoped to identify functional properties of traveling waves and test if they differ across brain regions.

Figure 3A,B illustrates the timecourse of mean directional consistency (DC) during the cue response interval for the traveling waves in the frontal lobe of Patient 26. DC indicates the tendency for the traveling waves on an electrode cluster to propagate in a single preferred direction. This plot indicates that the traveling waves on this cluster were not directionally organized at the moment of cue onset, but 500 ms later they reliably propagated anteriorly (DC=0.34). A different pattern was present for the traveling waves on an posterior electrode cluster in Patient 13 (Fig. 3C,D), whereby the directional organization was high at cue onset and subsequently decreased.

**Figure 3:**
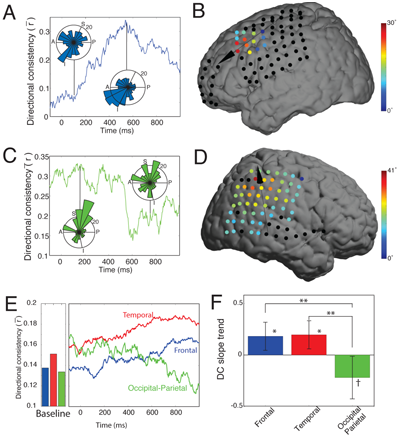
Temporal dynamics of traveling waves. (A) Timecourse of directional consistency (DC) for a traveling wave at 12.5 Hz from Patient 26’s frontal lobe. Inset circular histograms indicate the distributions of propagation directions across trials at the labeled timepoints. (B) Brain plot showing the mean relative phase shift at each electrode at the timepoint of peak consistency (see Panel B). (C & D) Traveling 6.2-Hz parietoccipital wave from Patient 13, which showed a decrease in DC after cue onset. (E) Timecourse of traveling-wave DC. Bars indicate the mean DC for each region when patient is out of task (F) Analysis of DC slope. Positive values indicate that DC increases following cue onset. Error bars denote 95% confidence intervals. Post-hoc test: ** denotes *p* < 0.01; *, *p* < 0.05; †, *p* < 0.1.

We confirmed that these patterns were reliable by measuring the timecourse of mean traveling wave DC at the group level. During the cue response interval, traveling waves in the temporal and frontal lobes showed increases in DC above baseline levels (Fig. 3E). Inversely, traveling waves from occipitoparietal clusters showed decreased DC during this same period. Thus, there are regional differences in the directional trends for traveling waves from different regions (Fig. 3F; ANOVA, *F*_(2,137)_ = 5.4, *p* < 0.01).

After a person views a stimulus, the brain exhibits stimulus-locked neural patterns, including phase resets of ongoing brain oscillations and new evoked neural signals (Rizzuto et al., 2003). Because these time-locked signals have theta- and alpha-band spectral components (Jacobs et al., 2006), we considered the possibility that they affected our measurements of traveling waves. We examined this issue by identifying time- (evoked) and phase-locked signals on electrode clusters with traveling waves. We the compared the timing of these signals to the timecourse of the traveling waves on the same channels (Fig. S4). Evoked signals and phase resets were prominent ∼200-400 ms post stimulus whereas traveling wave DC peaked later (∼800 ms). Furthermore, across clusters there was no correlation between the peak timepoint of traveling wave DC and the timepoints of the strongest evoked or phase-reset activity (*p*’s> 0.1). These results suggest that human cortical traveling waves we measured are not an artifact of stimulus-locked signals.

We next examined whether traveling waves correlated with the efficiency of memory processing. We compared the DC of the traveling waves on each electrode cluster between trials where patients had fast versus slow reaction times (median split). Overall, DC positively correlated with performance, such that traveling waves moved more reliably in the preferred direction for each cluster on trials when the subject responded quickly to the retieval cue (Fig. 4). This effect was significantly stronger for traveling waves in the frontal lobe (*F*_(2,137)_ = 4.5, *p* = 0.013), consistent with the notion that frontal theta oscillations are implicated in working memory (Jensen and Tesche, 2002; Onton et al., 2005).

**Figure 4:**
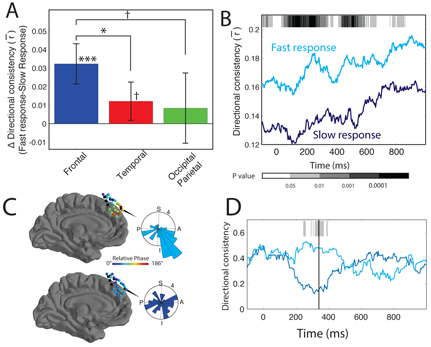
Traveling waves and behavior. (A) Mean difference in DC between fast and slow trials for 1 s after cue onset, separately calculated for each region. (B) Timecourse of mean DC in the frontal lobe between fast and slow trials. Gray shading indicates significance (paired *t* tests). (C) Time course of DC for data from Patient 3 that demonstrated elevated DC during trials where the patient responded rapidly. Shading indicates *p* values from a non-parametric circular direction distribution test (Fisher, 1993) between fast and slow response trials. (D) Brain plot showing the mean relative phase distribution across an oscillation cluster in patient 3. Inset plot shows distribution of propagation directions across trials 220 ms after probe onset. (E) Same as D, for trials where the patient performed badly. Post-hoc test: *** denotes *p* < 0.001; *, *p* < 0.05; †, *p* < 0.1.

We also examined how other properties of traveling waves varied in relation to memory efficiency (Table S3). On trials where patients had fast reaction times there was increased traveling wave PGD (*p* = 0.003) and oscillatory power (*p* < 0.001). There were no reliable performance-related changes in traveling waves’ temporal frequency, spatial frequency, or propagation speeds (all *p*’s> 0.5). These results indicate that efficient neural processing is predicted by traveling waves maintaining their optimal propagation direction rather than moving at a faster speed or oscillating at a higher temporal frequency.

### Mechanisms of cortical wave propagation

We next considered the neural mechanisms underlying traveling wave propagation. In animal model systems, identifying the mechanisms of traveling-wave propagation is an area of active research (Ermentrout and Kleinfeld, 2001; Sato et al., 2012). At first blush, examining this issue in humans might be even more challenging than in animals, because human brain oscillations are rather variable across time and frequency (Watrous et al., 2013) and because oscillations at neighboring frequencies in humans like alpha and theta are often considered to have different physiological roles (Roux and Uhlhaas, 2014). Nonetheless, we considered the possibility that a single physiological process could support traveling waves at multiple frequencies. We would be confident in identifying such a process if it fit the properties of wave propagation across the range of frequencies where we observed traveling waves. To understand the nature of wave propagation we first focused on one particular characteristic: propagation speed. We measured the propagation speed for each traveling wave on each individual trial (Figs. 5A & B, S5). Individual traveling waves showed a range of mean propagation speeds (mean=0.45 m/s, IQR 0.29–0.76), which indicated that this factor showed substantial variability for any model to explain.

**Figure 5:**
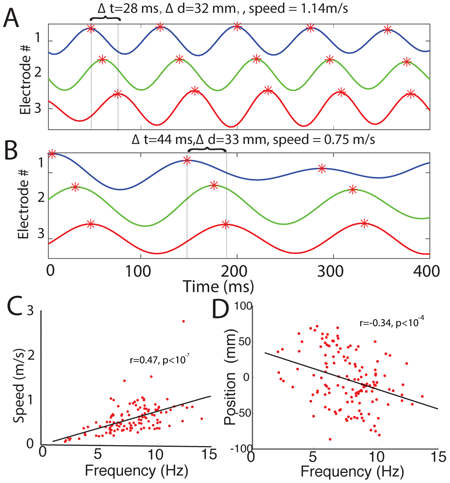
Characteristics of traveling-wave propagation. (A) Simultaneously recorded waveforms from three electrodes in a cluster with a 12.5-Hz traveling wave in Patient 26. This wave propagated at 1.14 m/s. Black dotted lines indicate the onset times of two oscillatory peaks. Labels indicate calculations used to measure this wave’s propagation speed; Δ*t*, delay between the arrival time of the oscillation between electrodes 1 & 3; Δ*d*, distance in mm between electrodes 1 & 3, after correction for the angle of incidence of wave propagation. (B) Plot showing the measurement propagation speed for a 7.5-Hz traveling wave from Patient 39, which moved at 0.75 m/s on this trial. (C) Population analysis of the relation between traveling-wave propagation speed and frequency. Each point indicates the mean frequency and propagation speed of traveling waves from one oscillation cluster. Black line is a least-squares fit. (D) Population analysis of the relation between traveling-wave frequency and cluster location along the anterior–posterior axis (Talairach coordinates [mm]).

Two theoretical models for traveling waves are the single-oscillator (SO) model and excitable-network (EN) models (Ermentrout and Kleinfeld, 2001). In the SO model, a traveling wave is produced when one oscillator projects to downstream areas via an array of gradually increasing conduction delays. In the EN model, traveling waves result from the gradual progression of excitatory activity across the cortex. Critically, the SO and EN models both predict that cortical traveling waves would move at a constant propagation speed, because the spatial propagation is a direct result of neuronal conduction delays, which are constant. In contrast to the prediction of these models, we found a positive correlation between propagation speed and oscillation frequency—waves with faster temporal frequencies propagated more rapidly, which seemingly rejects the EN and SO model. This frequency–speed correlation was present both at the group level, by comparing mean propagation speeds of waves from different electrode clusters (*r* = 0.47, *p* < 10^−7^; Fig. 5C), and at the single-trial level, by measuring separate waves observed on different timepoints from the same electrodes (Fig. S5A).

A third theoretical model is a network of weakly coupled oscillators (WCO) (Ermentrout and Kleinfeld, 2001), which have been used to model traveling waves in both neural and non-neural biological systems (Diamant and Bortoff, 1969; Ermentrout and Kopell, 1984). In the WCO model traveling waves are produced when there is a network of Kuramato oscillators (Kuramoto, 1981) that show two properties: First, the oscillators must be arranged in a linear array with weak phase coupling only between nearest neighbors. Second, there must be a spatial gradient in intrinsic frequency across the array of oscillators. When these two criteria are satisfied, oscillations propagate in the direction of the frequency gradient, towards oscillators with slower intrinsic frequencies (Ermentrout and Kopell, 1984). Critically, the WCO model predicts that oscillations with faster temporal frequencies propagate more rapidly because the traveling wave is a result of phase coupling (Ermentrout and Kopell, 1984; Ermentrout and Kleinfeld, 2001). Because, we found a positive correlation between propagation speed and oscillation frequency, as stated above, it indicates that human cortical traveling waves are driven by WCOs.

Furthermore, we found that the WCO model predicted the direction of wave propagation that we observed. In the WCO model, traveling waves propagate towards the oscillators with the slowest intrinsic frequencies (Ermentrout and Kopell, 1984). The human cortical traveling waves that we observed showed a correlation between propagation direction and the frequency of the local oscillation. At the group level, there was a systematic decrease in mean oscillation frequency along the brain’s posterior-to-anterior axis (*r* = 0.34, *p* < 0.001, Fig. 5D; Fig. 2A), which matches the mean propagation direction that also followed this same direction. This supports the idea that traveling waves generally propagate in a posterior-to-anterior direction because they are coordinated by an overall decrease in intrinsic oscillation frequency from posterior to anterior regions (Voytek et al., 2010).

The WCO model predicts that the propagation direction for each traveling wave is determined by the orientation of the local oscillation frequency gradient. We hypothesized that this could explain why while the majority of traveling waves propagated in a posterior-to-anterior direction, some exceptional oscillation clusters reliably showed traveling waves with the opposite propagation direction (e.g., Fig. 1). We thus compared the directions of frequency gradients and wave propagation at the individual cluster level to provide a more detailed test of the WCO model. Figures 6B & C show the direction distributions of traveling-wave propagation and of frequency gradients at the single-trial level for the oscillation cluster in Patient 1. Notably, both of these distributions have similar mean directions, suggesting that these factors are related. We next examined the correspondence between the directions of wave propagation and frequency gradients at the group level. Here, for each oscillation cluster with a traveling wave, we computed the direction of the mean frequency gradient and the mean direction of wave propagation and then computed their difference. The pairwise direction differences were clustered near zero (Rayleigh test *p* < 0.01; Fig. 6D), which suggests that there was a correlation between the direction of frequency gradients and propagation directions for individual clusters.

**Figure 6:**
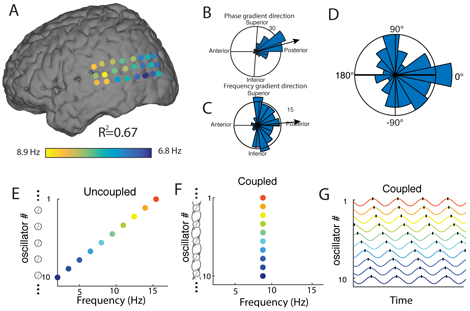
Mechanisms of traveling waves. (A) The instantaneous frequency distribution across electrodes of a traveling wave from same patient as Fig. 1 on one trial, demonstrating an anterior-to-posterior decreasing spatial-frequency gradient (*r*^2^ = 0.67). (B) Distribution of the direction of traveling-wave phase propagation across all trials on this cluster (reproduced from Figure 1F). (C) Distribution of the direction of the spatial frequency gradients across these electrodes. In B & C, black lines indicate the mean gradient directions, thus demonstrating a correspondence between the directions of phase and frequency gradients. (D) Distribution of angular differences between each traveling-wave’s mean phase gradient direction (propagation direction) and frequency gradient direction. The mean of the distribution is clustered near zero degree suggesting the propagation of each wave matches with their own frequency gradient (*p* < 0.01, rayleigh test). (E–G) Illustration of a model of weakly coupled oscillators (Ermentrout and Kopell, 1984) with parameters matched to our findings. Color indicates intrinsic frequency. When there is no phase coupling (Panel E), individual oscillators demonstrate their intrinsic oscillation frequencies from 2 Hz (anterior) to 16 Hz (posterior). When phase coupling is present (Panels F–G), all oscillators have the same temporal frequency (F) and a traveling wave emerges (G).

In summary, the WCO model provides a good fit for our results because it illustrates how the direction and speed of the traveling waves we observed can be predicted by the local frequencies of the oscillations we measured. The WCO model suggests that the existence and direction of traveling waves are modulated by two factors: the strength of local phase coupling and spatial gradients of intrinsic oscillation frequencies (Ermentrout and Kopell, 1984; Ermentrout and Kleinfeld, 2001). When phase coupling is absent, there are no traveling waves because intrinsic oscillation frequencies differ between electrodes (Fig. 6E). When phase coupling is present, traveling waves emerge, propagating in the direction of decreasing oscillation frequency (Fig. 6F-G).

## Discussion

Our findings demonstrate a new functional role for theta- and alpha-band oscillations by behaving as traveling waves, in addition to their roles supporting neuronal coding (O’Keefe and Recce, 1993), modulating synaptic plasticity (Huerta and Lisman, 1995), and various other functional processes (Klimesch, 1999; Buzsáki and Draguhn, 2004). Researchers had previously known that oscillations modulated cortical interactions in a point-to-point fashion between specific cortical areas (Hyman et al., 2011; Liebe et al., 2012). Our results add to this work by demonstrating that oscillations also coordinate neural processing across large contiguous extents of human cortex. The theta and alpha bands that displayed traveling waves is also known to participate in the phenomenon of cross-frequency coupling (CFC; Canolty et al. 2006). In CFC, the local oscillatory phase is coupled to the simultaneous amplitude of neuronal spiking (Jacobs et al., 2007) and to high-frequency oscillations (Canolty et al., 2006; Bahramisharif et al., 2013). If individual regions simultaneously exhibit both traveling waves as well as CFC (Bahramisharif et al., 2013), it indicates that bands of neuronal spiking activity propagate across the cortex during cognition. Based on the measurements of the traveling waves we observed, it suggests that these bands have a width of ∼2.4–4.8 cm and a mean propagation speed of ∼45 cm/s.

Our findings suggest that human cortical traveling waves can be modeled as a network of weakly coupled oscillators (WCO). In addition to suggesting a mechanism underlying traveling waves, the WCO model has practical implications for understanding traveling wave dynamics in behavior. A key part of the WCO model is the existence of a link between local intrinsic oscillation frequency and the direction of signal propagation. Traditionally, the frequency of a brain oscillation has been considered to be important because it indicates the functional role of a given oscillation. For example, oscillations in the neighboring theta and alpha bands have been associated with distinct functions (memory and idling, respectively; Klimesch, 1999; Roux and Uhlhaas, 2014). Instead, our results suggest that the frequency of a human cortical oscillation does not link to a specific function but instead that it indicates the regions where signals are propagating. Thus, based on the WCO model, our results provide a new way for predicting the direction of neuronal information flow by measuring the local oscillation frequency. Furthermore, our findings and the WCO model suggest a way for distinguishing periods of decreased cortical connectivity because they would be indicated as periods when neighboring cortical regions exhibit frequency gradients rather than traveling waves.

Studies of perception find that human sensory processing is segmented into discrete behavioral states. Scalp recordings indicated that these individual states correspond to separate cycles of alpha oscillations (VanRullen and Koch, 2003). Combined with our findings, it suggests that individual traveling-wave cycles represent spatially discrete pulses of neural activity that correspond to distinct behavioral states. This suggests that traveling waves could be relevant functionally for large-scale computation by propagating discrete “packets” of neuronal activity that each correspond to an individual state (Freeman, 2003). In this way, cortical traveling waves may provide a timing mechanism to distinguish distinct network states across space and time, in a similar fashion to theta oscillations in the hippocampus, where individual oscillation cycles temporally segment the representations of different cognitive maps (Jezek et al., 2011). Synchronized oscillations in multiple regions can form carrier signals that allow the representation of detailed behavioral information via phase and amplitude modulation (Freeman and Schneider, 1982; Agarwal et al., 2014)—it is possible that neocortical traveling waves are relevant for this phenomenon by propagating these complex oscillatory representations (Freeman, 2003).

Across the brain, most traveling waves propagated in a posterior-to-anterior direction, which suggests that they play a role in communicating information from sensory regions to high-level frontal brain regions. Posterior-to-anterior traveling waves may also support detailed features of neural computations, as many brain areas also show posterior-to-anterior gradients in the abtractness of their neural representations, such as the coding of spatial location in the hippocampus (Kjelstrup et al., 2008), the representation of task rules in the frontal lobe (Badre and D'Esposito, 2009), and the nature of object representations in the temporal lobe (Harry et al., 2016). Although they were in the minority, some traveling waves propagated in an anterior-to-posterior direction. In the monkey visual system, alpha-band oscillations travel from high-level (anterior) to low-level (posterior) regions and have been linked functionally to top-down feedback processes (Bastos et al., 2015). Likewise, in humans it is possible that the function of traveling waves in a given region varies according to its direction.

Some elements of the planar traveling waves we observed were noted in previous studies that used other methods to examine spatial characteristics of human brain signals (Patten et al., 2012; Bahramisharif et al., 2013; Alexander et al., 2013). A differentiating feature of our approach was that we identified traveling waves across multiple regions, directions, and frequencies directly in individual subjects and trials. Because individual subjects exhibited traveling waves at different frequencies and directions in many regions, it suggests that there are true intersubject differences in the spatial and temporal structure of brain oscillations and traveling waves that are not possible to accurately capture with group-average analyses. In addition to traveling waves, the brain also exhibits other large-scale spatial patterns of oscillations, such as phase cones and spirals (Freeman and Barrie, 2000; Muller et al., 2016), as well as smaller patterns at finer spatial scales (Freeman, 2003; Rubino et al., 2006). Together, this work suggests that there is potential for researchers and engineers to identify important new spatial patterns of brain dynamics across the cortical surface using improved high-resolution electrocorticographic electrodes (Viventi et al., 2011; Khodagholy et al., 2015) rather than necessarily requiring penetrating electrodes or single-cell recordings.

Although our data came from epilepsy patients, there are several reasons why we believe our results generalize to healthy humans. Our data analysis focused on theta- and alpha-band oscillations in a memory task, which are known to be engaged similarly in healthy subjects. Brain recordings from normal college students show the same types of task-related theta and alpha changes that we observed here (Jacobs et al., 2006). Furthermore, although epileptic activity often shows spatial propagation across the cortex (e.g., Liou et al., 2017), the spatial and temporal characteristics of interictal and seizure-related activity dif er compared to the traveling waves we described, suggesting that these are distinct phenomena. Finally, analyses of noninvasive brain recordings from healthy individuals have reported spatial patterns that are consistent with the traveling waves we observed (Patten et al., 2012; VanRullen and Lozano-Soldevilla, 2017).

In addition to demonstrating a new fundamental feature of human brain activity, the ability to measure traveling waves noninvasively may have significant practical implications. Because our results show that neuronal oscillations can be synchronized across large regions of cortex, researchers and clinicians examining noninvasive brain recordings should consider that aspects of their findings may result from large neural masses (Freeman, 2003) rather than precisely localizable point sources (Michel et al., 2004). Furthermore, whereas many electrical signals from the brain are commonly interpreted as event-related potentials (ERPs) or as task-induced power changes from local oscillators, instead it is possible that these signals could result from traveling waves that become transiently organized at a particular timepoint and phase across a cortical region (Alexander et al., 2013). The potential for non-invasively measuring traveling waves on a single-trial basis may be useful for the development of brain–computer interfaces (BCI). However, for traveling waves to be useful for BCIs, given the intersubject differences we observed (Fig. 1), it seems important to characterize these patterns individually for each subject rather than averaging across individuals. In this way, measuring traveling waves' instantaneous properties may provide a new tool for neural interfacing, by tracking a subject’s attention or cognitive state for timing stimulus presentation or neuromodulation (Ezzyat et al., 2017).

In summary, our findings show that traveling waves of neuronal oscillations comprise large spatiotemporal patterns across the human cortex (Livanov, 1977; Freeman, 2003). The existence of traveling waves indicates that neuronal oscillations with specific frequency characteristics are important for large-scale brain connectivity. We hypothesize that traveling-waves relate to the slower functional connectivity signals that have been identified with fMRI (Honey et al., 2007), based on the known link between fMRI activity and the power of neuronal oscillations (Debener et al., 2005). Thus, traveling waves expand our understanding of cortical functional connectivity by showing that signal propagation across neural networks can be transient and rhythmic. Our findings emphasize that human cognition is supported by complex, large-scale neural patterns that are exquisitely organized across both time and space. Traveling waves may be important for understanding the nature of large-scale brain connectivity in cognition by showing when behavioral information is represented and where it is propagating.

## Methods

### Subjects and Task

We examined direct brain recordings from 77 epilepsy patients who had electrodes surgically implanted to guide seizure mapping. Individual patients were implanted with a configuration of electrodes customized according to their clinical needs, which included both electrocorticographic (ECoG) surface grid and strips as well as depth electrodes. Our data collection was a continuation of previous reported study (Jacobs and Kahana, 2009) and recordings were made at four hospitals (Thomas Jefferson University Hospital, Philadelphia; University of Pennsylvania Hospital Philadelphia; Children’s Hospital of Philadelphia, and University Hospital Freiburg). All patients consented to having their brain recordings used for research purposes and the research was approved by relevant Institutional Review Boards. For the work described here, we examined only ECoG grid and strip electrodes on the cortical surface. The spacing between neighboring electrodes was 10 mm (center-to-center).

During free time between clinical procedures, these patients performed the Sternberg working memory task (Sternberg, 1966) on a laptop computer at their bedside. Each trial of the task consisted of two phases. In the first phase they memorize a short list of items. The second phase involves memory retrieval. Here they view a probe item and press a key to indicate if the probe was present in the remembered list. Task performance was excellent, with patients having a mean accuracy of 90% and mean reaction time of 1.76 s. Our data analyses examined brain recordings during memory retrieval because it let us compare properties of patients' brain signals to their simultaneous behavioral performance (Jacobs et al., 2006).

### Data acquisition

The electrical activity from each electrode was recorded by a clinical recording system whose timing was synchronized with the task computer. We pre-processed the data by downsampling the recordings to 250 Hz and performing an anatomically weighted average re-referencing (Jacobs and Kahana, 2009). We identified the location of each recording electrode by co-registering a pre-surgical structural magnetic resonance image (MRI) image with a post-operative computed tomography (CT) image. From these images, we identified the location of each recording contact on the CT images and computed the electrode location in standardized Talairach coordinates (Talairach and Tournoux, 1988).

### Identifying spatial clusters of electrodes with similar oscillations

Given our interest in characterizing propagating traveling waves, we designed an algorithm to identify spatial clusters of electrodes with narrowband oscillations at the same frequency. This algorithm accounted for several complexities of human brain oscillations measured with ECoG signals, including differences in electrode positions across subjects and variations in oscillation frequencies across individuals (Fig.S2).

In this procedure, first we used Morlet wavelets (wave number 6) to compute the power of the neuronal oscillations throughout the task at 129 frequencies logarithmically spaced from 2 to 32 Hz. To identify narrowband oscillations at each site, we fit a line to each patient’s mean power spectrum in log-log coordinates using a robust linear regression (Fig. S1; Manning et al. 2009; Lega et al. 2012; Zhang and Jacobs 2015). We then subtracted the actual power spectrum from the regression line. This normalized power spectrum provides a “whitened” version of the signal that removes the 1/*f* background signal and emphasizes narrowband oscillations as positive deflections. We identified narrowband peaks in the normalized power spectrum as any local maximum greater than one standard deviation above the mean (Zhang and Jacobs, 2015).

Next, we implemented a spatial clustering algorithm to identify oscillation clusters, which we defined as contiguous groups of electrodes in each subject that exhibited narrowband oscillations at the same frequency. First, considering a series of 2-Hz-wide intervals centered at 2 to 32 Hz in 1-Hz steps, we identified all the electrodes that exhibited a narrow-band oscillatory peak within that range. We counted the number of electrodes with oscillatory peaks at each frequency, and computed local maxima as potential oscillation clusters. We then tested whether the electrodes that contributed to each local maximum comprised a spatially contiguous group. We found all the electrodes with a peak in the 2-Hz interval around each local maximum and created a pairwise-adjacency matrix to judge their spatial proximity. This matrix indicated whether each electrode pair was separated by less than 15 Talairach units (15 mm). Finally, we used this adjacency matrix to identify mutually connected spatial clusters of electrodes by computing the connected components of this graph using Tarjan’s (1972) algorithm. We included in our subsequent analyses only clusters with at least four electrodes.

A key feature of our spatial clustering algorithm is that it adapts to the specific anatomical orientation of each patient’s ECoG electrode organization by utilizing Talairach coordinates rather than labeled positions from clinical recordings. Thus, our methods are capable of identifying oscillation clusters that span multiple ECoG grids or strips (e.g., Fig. 1H), rather than being limited to identifying signals within regularly structured electrode arrays (Rubino et al., 2006; Lubenov and Siapas, 2009; Zhang and Jacobs, 2015).

### Identifying traveling waves

Having identified groups of electrodes with oscillations at the same frequency, we next sought to identify traveling waves. Intuitively, a traveling wave can be described as an oscillation that moves progressively across a region of cortex. Quantitatively, a traveling wave can be described as a set of simultaneously recorded neural oscillations at the same frequency whose instantaneous phases vary systematically with the location of the recording electrode. Although many types of traveling waves are possible (Ermentrout and Kleinfeld, 2001; Muller et al., 2016), we focused our analyses here on one specific form of this phenomenon, the planar wave (Lubenov and Siapas, 2009). A planar wave would appear in our dataset as an oscillation cluster whose instantaneous phases exhibited a linear relationship with electrode location.

We followed the following procedure to identify planar waves in the phases from each oscillation cluster. First, we measured the instantaneous phases of the signals across each oscillation cluster by applying a Butterworth filter to the signals from each electrode at the cluster’s narrowband peak frequency (3-Hz bandwidth). Then we perform the Hilbert transform on each electrode’s filtered signal to extract the instantaneous phase at each timepoint (Freeman, 2007). To facilitate visualization, we rotate the phase distributions so that the smallest value is set to 0.

We used circular statistics to identify planar waves of phase progression across each oscillation cluster at each timepoint (Fisher, 1993). For each spatial phase distribution, we used a two-dimensional circular-linear regression to assess whether the observed phase pattern varied linearly with the electrode’s coordinates in 2-D. In this regression, for electrode *i*, *x_i_* and *y_i_* represent the 2-D coordinates and *θ_i_* is the instantaneous phase. *x* and *y* are determined by projecting the 3-D Talairach coordinates for each cluster into the best-fitting 2-D plane.

A 2-D circular-linear model has three parameters to be fit: the phase slopes *a* and *b*, which each correspond to the rate of phase change (or spatial frequencies) in each dimension, and the phase offset 0. We converted this model to polar coordinates to simplify fitting. This polar model has two parameters: The angle of wave propagation *α*, defined as *α* = atan2(*b, a*), and the spatial frequency *ξ*, defined as 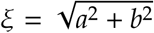. Circular-linear models do not have an analytical solution and must be fitted iteratively (Fisher, 1993). We fit *α* and *ξ* to the distribution of oscillation phases at each timepoint by conducting a grid search over *α* ∈ [0°, 360°] and *ξ* ∈ [0,18] in increments of 5°and 0. 5°/mm, respectively. (Note that ξ = 18 corresponds to the spatial Nyquist frequency of 18°/mm.) The model parameters for each timepoint are fitted to most closely match the phase observed at each electrode in the electrode cluster. For each value of *α* and *ξ*, the model’s predicts the phase 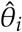 at each electrode *i* as

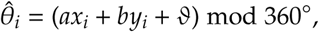

where *α* = *ξ* cos(*α*) and *b* = *ξ* sin(*α*). Then we compute the goodness of fit as the mean vector length *r*̅ of the residuals between the predicted 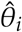 and actual (*θ_i_*) phases (Fisher, 1993),

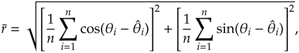

The selected values of *α* and *ξ* are chosen to maximize *r*̅.

Figure S3 illustrates the results of performing this procedure for several trials from Patient 1. The fitted model coefficients indicate physical characteristics of any identified planar traveling wave, by showing the slope and direction of the spatial phase gradient: *α* is the wave’s instantaneous propagation direction and *ξ* is it’s spatial frequency (i.e., rate of phase change over space) in °/mm.

To measure the statistical reliability of each fitted traveling wave we examined the phase variance that was explained by the best fitting model. As in earlier work (Kempter et al., 2012), we adapted the *r*^2^ goodness-of-fit measure from linear correlation for use with circular data. To do this, we computed the circular correlation *ρ_cc_* between the predicted 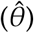 and actual (*θ*) phases at each electrode (Kempter et al., 2012):

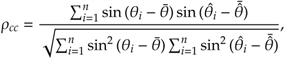

where bar denote averaging across electrodes. We squared the result to compute 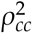 Finally, we applied an adjustment to account for the number of fitted model parameters:

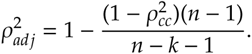

where *n* is the number of electrodes, and *k* is number of independent regressors (k=3). We refer to 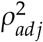 as *phase gradient directionality* (PGD) in the main text, similar to Rubino et al. (2006).

### Classification of traveling waves

The procedure described above identifies the spatial gradients in a single instantaneous phase distribution. Next, we describe the procedure we used to measure the reliability of each phase gradient in terms of PGD and directional consistency. First, we computed the mean PGD for each trial and cluster. We computed the median PGD for each cluster (across trials) and compared this value with the distribution of PGDs estimated from a shuffling procedure (1,000 iterations). In this shuffling procedure, on each trial we compute a single PGD value after randomly permuting the locations of individual electrodes within each cluster, while preserving other features of the data. We identify reliable traveling waves as electrode clusters in which the actual PGD value exceeds 95% of the distribution of PGD values from shuffling. Due to the spatial permuting that occurs in the shuffling, this method ensures that the traveling waves we identify reliably exhibit planar waves compared to other types of phase patterns with less structure (Maris et al., 2016).

Finally, here we considered only the electrode clusters with traveling waves that propagated in a consistent direction over time. To do this, after identifying the clusters with reliable PGD values, we computed for each cluster the distribution of propagation directions (*α*) across trials. The clusters we designate as exhibiting significant traveling waves are the ones that exhibit a non-uniform distribution of propagation directions, as determined by the Rayleigh test (Fisher, 1993) at *p* < 0.05. We calculated the directional consistency (DC) of each cluster as the circular mean vector length of this distribution of propagation directions. Thus, a cluster with DC=1 indicates that its traveling waves always propagate in the same direction.

### Statistical assessment of traveling-wave properties

We performed a series of analyses to compare the properties of traveling waves between clusters from different brain regions. To compare directional propagation between different traveling waves, we first converted each cluster’s mean propagation direction into Talairach coordinates. Note that this sometimes caused directional plots to appear distorted, when the 2-D plane for each cluster was skewed relative to the Talairach axes (e.g., Fig. 1F). Then, to assess regional differences in propagation directions (Fig. 2B), we grouped each cluster according to whether it was in the frontal, temporal or occipitoparietal region. When a cluster spanned multiple areas, we labeled it to the lobe that contained the plurality of its electrodes.

To measure the temporal frequency *f* of a traveling wave, we followed *f* = *dθ*̅/*dt*, where *θ*̅ is the average phase at each timepoint, taken as the circular mean across electrodes, and *t* is time. We used this approach to assess each wave’s propagation speed *v* as *v* = *f*/*ξ*, where *f* is instantaneous temporal frequency and *ξ* is the spatial frequency. To summarize the propagation speed for each wave (Fig. 5C), we computed the median propagation speed across all timepoints, excluding periods where the model fit was poor (i.e., mean PGD < 0.5; Rubino et al., 2006). Twenty one clusters had no timepoints with mean PGD ≥ 0.5 and thus were excluded from this analysis. To identify task-related changes in wave DC, we computed the trend of DC over time for each traveling wave cluster (Fig. 3F) as the Pearson correlation between mean DC and time.

We considered the possibility that traveling waves could relate to stimulus-locked signals by calculating, for each cluster that showed a traveling waves, the mean phase resetting *R*̅ and event-related potential (ERP) (Rizzuto et al., 2003). To identify the contribution of phase resetting to each cluster’s post-stimulus activity, we calculated the phase distribution at each point in time for the cluster’s mean frequency as in Rizzuto et al. (2003). We calculated the ERP for each cluster by first filtering the raw signal at 0.5–40 Hz, performing –200–0 baseline correction, and finally identifying the individual electrode with the largest absolute ERP component.

To assess the role of traveling waves in behavior, for each cluster we compared the properties of traveling waves between trials where the patient had fast or slow responses to the probe in the memory task (median split). Then, for each cluster we computed the mean values of several properties of traveling waves separately for fast and slow trials: power, temporal frequency, PGD, spatial frequency, and directional consistency (Tab. S3). Unless otherwise specified, we used Student’s *t*-tests and ANOVAs to compare differences between regions, with rejection of the null hypothesis reported after FDR correction (Genovese et al., 2002) at *q* = 0.05.

### Model of traveling waves based on weakly coupled oscillators

A linear array of weakly coupled Kuramoto oscillators (Kuramoto, 1981) produces traveling waves when it exhibits two properties: (1) weak phase coupling only between neighboring oscillators and (2) a spatial frequency gradient. In this spatial frequency gradient, the intrinsic frequency of each oscillator systematically varies with position along the array (Ermentrout and Kopell, 1984; Ermentrout and Kleinfeld, 2001). This configuration explains the appearance of traveling waves in the digestive system (Diamant and Bortoff, 1969) and is hypothesized to generate traveling waves in the cortex (Ermentrout and Kleinfeld, 2001).

We simulated this model to demonstrate that the frequency gradient we observed in our data (Fig. 6A) is feasible for producing traveling waves that propagate in the directions we observed (Fig. 1). As in earlier work (Ermentrout and Kleinfeld, 2001), we implemented the Kuramoto model as

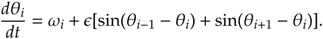

where *i* indexes separate oscillators, *ω_i_* is an oscillator’s intrinsic frequency, *θ_i_* is the instantaneous phase, and *є* is the strength of phase coupling between neighboring oscillators. We used this model to simulate ten oscillators, which vary in frequency from 2 Hz to 16 Hz (*ω_i_* = 1.56*i* + 0.44 for *i* ∈ [1,10]), corresponding to a decreasing posterior-to-anterior frequency gradient from oscillator 10 to 1. Our simulations showed that when there is no cortico-cortico coupling (*є* = 0), each oscillator exhibits independent oscillations (Fig. 6E)—this resembles the spatial frequency gradients we observed in individual subjects (Fig. S6) and at the population level (Fig. 5D). When coupling is positive (*є* ≫ 0), the same distribution of intrinsic frequencies produces a traveling wave propagating in a posterior-to-anterior direction (Fig. 6F–G).

## Acknowledgements

J.J. acknowledges support from NIH Grant MH104606. We wish to thank Jonathan Miller, Salman Qasim, and Melina Tsitsiklis for technical discussions, and Michael Kahana for help with data collection. Raw data for this paper are available at http://memory.psych.upenn.edu/.

## Author Contributions

J.J. conducted the experiments. H.Z., J.J., A.P., and A.J.W. analyzed the data. J.J., A.J.W., and H.Z. wrote the paper.

